# A universal, open-source, high-performance tool for automated sleep staging

**DOI:** 10.1101/2021.05.28.446165

**Authors:** Raphael Vallat, Matthew P. Walker

## Abstract

The creation of a completely automated sleep-scoring system that is highly accurate, flexible, well validated, free and simple to use by anyone has yet to be accomplished. In part, this is due to the difficulty of use of existing algorithms, algorithms having been trained on too small samples, and paywall demotivation. Here we describe a novel algorithm trained and validated on +27,000 hours of polysomnographic sleep recordings across heterogeneous populations around the world. This tool offers high sleep-staging accuracy matching or exceeding human accuracy and interscorer agreement no matter the population kind. The software is easy to use, computationally low-demanding, open source, and free. Such software has the potential to facilitate broad adoption of automated sleep staging with the hope of becoming an industry standard.

## Introduction

Sleep is fundamental to human health. Adequate sleep supports numerous physiological body systems, including immune, metabolic and cardiovascular function (Besedovsky et al., 2019; Cappuccio and Miller, 2017; Harding et al., 2020). For the brain, sufficient sleep facilitates optimal learning, memory, attention, mood and decision making processes (Ben Simon et al., 2020; Walker, 2009). Sleep has emerged as a preventive strategy to reduce the risk of cardiovascular and metabolic disease, the accumulation of Alzheimer’s disease brain pathology (Cappuccio and Miller, 2017;Ju et al., 2013; Winer et al., 2020) as well as all-cause mortality (Cappuccio et al., 2010; Leary et al., 2020).

Considering this impact, the demand for quantifying human sleep at a research-, clinical- and consumer-based level has increased exponentially over the past decade (Fleming et al., 2015; Shelgikar et al., 2016). However, and unlike the increasing automaticity of other health-based analysis methods, such as blood assays, areas of radiological medicine and genetics (Choy et al., 2018), there is no similar industry-standard, fully automated sleep-scoring system that is highly accurate, freely available, extensively validated on heterogeneous datasets, and easy to use no matter what level of technical expertise (Fiorillo et al., 2019).

Polysomnography (PSG) — the simultaneous measurement of brainwaves, eye movements, muscle activity, heart rate and respiration — is the gold-standard for objective quantification of sleep. The classification of sleep stages across the nights provides information on the overall architecture of sleep across the night, as well as the duration and proportion of the sleep stages, both of which inform of potential sleep disorders or diseases. Currently, such sleep scoring is typically performed by humans, accomplished by first dividing the PSG recording into 30-seconds segments (called “epochs”). Each epoch is then assigned a sleep stage based on standard rules defined by the American Academy of Sleep Medicine (AASM, (Berry et al., 2012; Iber et al., 2007)).

Every night, thousands of hours of sleep are recorded in research, clinical and commercial ventures across the globe. However, since sleep staging is performed visually by human experts, it represents a pragmatic bottleneck. It is a non-trivial, time-intensive process, with visual scoring of a single night of human sleep typically requiring up to 2 hours to complete by a well-trained individual. As if not more critical, this human-scored approach suffers from issues of low *inter*-scorer consistency of agreement (~83% agreement, (Rosenberg and Van Hout, 2013)). Moreover, the same individual typically experiences low *intra*-scorer agreement of the same sleep recording (~90%, (Fiorillo et al., 2019)). That is, different human sleep-scoring experts presented with the same recording will likely end up with dissimilar sleep-staging evaluations, and even the same expert presented with the same recording at two different time points will arrive at somewhat different results.

Advances in machine-learning have led efforts to classify sleep with automated systems (for a review see (Fiorillo et al., 2019)). However, accurate automated sleep staging has not yet become a de facto standard in the field, despite its numerous advantages. There are several reasons for this. First, some algorithms are not free, sitting behind a paywall (e.g. pay-per-use license (Patanaik et al., 2018)). Second, other algorithms require paid software to run the algorithm on, such as MATLAB (Lajnef et al., 2015). Third, some algorithms have been trained on a sample size too small for robustness (e.g., less than 30 nights, (Zhang and Wu, 2018)). As a result, their ability to generalize to other recording systems and/or populations, including patients with sleep disorders or across broad age ranges, have been problematic. Fourth, algorithms are often too complicated for use by most individuals as they require moderate-to high-level programing experience, creating a barrier of entry (e.g., they require the users to manually train the classifier (Sors et al., 2018)), or are based on “black-box” algorithms (Malafeev et al., 2018; Phan et al., 2019).

Seeking to address these issues, here, we describe a free, flexible and easy to use automated sleep-staging software that has been trained and validated on more than 27,000 hours of PSG-staged sleep from numerous independent and heterogeneous datasets with a wide range of age, ethnicities, and health status levels. Furthermore, the algorithm offers a high level of sensitivity, specificity and accuracy matching or exceeding that of typical interscorer agreement no matter the characteristics of the sampled individuals (age, sex, BMI, sleep-disorder status). Finally, the algorithm and software is expressly designed to be flexible and elemental to use by all individuals, no matter what level of technical expertise, and from a computational perspective, requires nothing more than a basic entry-level laptop for full function.

## Methods

### Datasets

The algorithm was trained and validated on a collection of large-scale independent datasets from the National Sleep Research Resource (NSRR; https://sleepdata.org/; (Dean et al., 2016; Zhang et al., 2018) — a web portal funded by the National Heart, Lung and Blood Institute (NHLBI). This database offers access to large collections of de-identified physiological signals and clinical data collected in research cohorts and clinical trials. All data were collected as part of research protocols that were approved by the local institutional review board at each institution, with written and informed consent obtained from each individual before participation. All PSG recordings were scored by trained technicians using standard AASM guidelines (Iber et al., 2007; Silber et al., 2007). A full description of the datasets can be found on https://sleepdata.org/. The following datasets were used: 1) the Multi-Ethnic Study of Atherosclerosis (MESA), 2) the Cleveland Family Study (CFS), 3) the Cleveland’s Children’s Sleep and Health Study (CCSHS), 4) the Sleep Heart Health Study (SHHS), 5) the Osteoporotic Fractures in Men Study (MrOS), 6) the Childhood Adenotonsillectomy Trial (CHAT) and 7) the Home Positive Airway Pressure (HomePAP).

Each dataset was randomly split into training (up to 600 nights) and testing (up to 100 nights) sets. PSG nights included in the training set were used for model building and training, while PSG nights included in the testing set were used for performance evaluation. Importantly, the training and testing sets were completely separate (i.e. no overlap). The code used to generate the training and testing sets can be found <u>here</u>.

To provide an unbiased evaluation of the model on a completely new dataset, we further tested the performance of the algorithm on the Dreem Open Dataset (DOD, (Guillot et al., 2020)), a publicly-available dataset including healthy individuals and patients with sleep apnea. No nights from DOD were used for model training. Each night of the DOD was scored by 5 clinical experts, thus allowing to compare the performance of the algorithm against a consensus of human scorers (see **Consensus Scoring** section).

### Preprocessing and features extraction

The following five standardized preprocessing steps were applied to the datasets: 1) nights with poor PSG data quality and/or unreliable scoring were excluded (these variables are provided for each night on the NSRR website, 2) nights for which the human scoring did not include all sleep stages (e.g. REM or N3 sleep was missing) were excluded, 3) nights were cropped to 15 minutes before and after sleep to remove irrelevant extra periods of wakefulness or artefacts on both ends of the recording, 4) nights with a duration outside the range of 4 to 12 hours were also excluded, and 5) for each remaining night, we extracted a single central EEG, left EOG and chin EMG. We chose a central EEG (e.g. C4-M1 or C4-Fpz) since the American Association of Sleep Medicine (AASM) recommends that a central EEG should be included in all PSG recordings, and it is therefore more likely to be present in a variety of PSG recordings. These signals were then downsampled to 100 Hz to speed up computation time, and bandpass-filtered between 0.40 Hz and 30 Hz.

The classification algorithm is based on a machine-learning approach in which a set of “features” is extracted from the EEG signal, and optionally though non-necessarily, from the EOG and EMG signals as well. Consistent with human sleep staging, features are calculated for each 30-seconds epoch of raw data. An overview of the features implemented in the algorithm is provided below, focusing on the EEG features (although the EOG and EMG features are virtually identical). All code used to compute these features is made open-source and freely available to all (see **Data and code availability**). A full list of the features calculated by the algorithm can be found <u>here</u>.

#### Time-domain features

The implemented time-domain features comprise standard descriptive statistics, i.e., the standard deviation, interquartile range, skewness and kurtosis of the signal. In addition, several non-linear features are calculated, including: the number of zero-crossings, the Hjorth parameters of mobility and complexity (Hjorth, 1970), the permutation entropy (Bandt and Pompe, 2002; Lajnef et al., 2015), and the fractal dimension (Esteller et al., 2001; Higuchi, 1988; Petrosian, 1995) of the signal.

#### Frequency-domain features

Frequency-domains features were calculated from the periodogram of the signal, calculated for each 30-seconds epoch using Welch’s method (Welch, 1967) (Hamming window of 5 seconds with a 50% overlap [= 0.20 Hz resolution], median-averaging to limit the influence of artefacts). Features included: the relative spectral power in specific bands (slow = 0.4-1 Hz, delta = 1-4 Hz, theta = 4-8Hz, alpha = 8-12Hz, sigma = 12-16Hz, beta = 16-30 Hz), the absolute power of the broadband signal, as well as power ratios (delta / theta, delta / sigma, delta / beta, alpha / theta)

#### Smoothing and normalization

When scoring sleep, human experts frequently rely on contextual information, such as prior and future epochs around the current epoch being scored (i.e., what was the predominant sleep stage in the last few minutes, what stage is the next epoch). By contrast, feature-based algorithms commonly process one epoch at a time, independently of the past and future epochs, overlooking such contextual temporal information. To overcome this limitation, the current algorithm implemented a smoothing approach across all the aforementioned features. Specifically, the features were smoothed using two different rolling windows: 1) a 5.5 minutes centered and triangular-weighted rolling average, and 2) a rolling average of the last 5 minutes prior to the current epoch. In other words, this allows to integrate some data from the past and future into the current epoch. The use of a ~5 min temporal window offers an optimal compromise between short-term and long-term sleep dynamics.

Critically, there is marked natural inter-individual variability in EEG brainwave activity (Buckelmüller et al., 2006; De Gennaro et al., 2008), meaning that each individual has a unique EEG fingerprint. To take this into account, a subset of features were z-scored across each night, i.e. expressed as a deviation from the night’s average. The inclusion of such normalized features aids in accommodating the error-potential impact of inter-individual variability upon the algorithm, and thus improves final accuracy.

Finally, the features set includes time elapsed from the beginning of the night, normalized from 0 to 1. This importantly accounts for the known asymmetry of sleep stages across the night i.e., the predominance of deep NREM sleep in the first half of the night, and conversely, a preponderance of REM and lighter NREM sleep in the second half of the night.

If desired, the user can also add information about the participant’s characteristics, such as the age and sex that are known to influence sleep stages (see **Results**, (Carrier et al., 2001; Ohayon et al., 2004)), which the classifier then takes into account during the staging process.

### Machine-learning classification

The full training dataset was then fitted with a LightGBM classifier (Ke et al., 2017) — a tree-based gradient-boosting classifier using the following hyper-parameters: 300 estimators, maximum tree depth of 7, maximum number of leaves per tree of 70 and a fraction of 80% of all features selected at random when building each tree. All of these parameters were chosen to prevent overfitting of the classifier to the training set i.e., a higher tree depth could result in better accuracy, but this comes at the cost of a lower generalizability of the model.

In addition, custom sleep stage weights were passed to the classifier to limit the imbalance in the proportion of sleep stages across the night. Without such weighting, a classifier would favor the most represented sleep stage (N2, ~50% of a typical night), and conversely, would seldom choose the least-represented sleep stage (N1, ~5%). The best weights were found by running a 2-fold cross-validation grid search on a subsample of the training nights (25% of each dataset), with accuracy and F1-macro as optimization metrics. The resulting optimal weights were as follows: 2 for N1,1 for N2 and Wake, and 1.2 for N3 and REM. The pre-trained classifier was then exported as a compressed file (~2 MB) and used to predict 1) a full hypnogram for each night of the testing set, 2) the associated probabilities of each sleep stage for each 30-seconds epoch.

### Performance evaluation

Performance of the algorithm was evaluated using established standardized guidelines (Menghini et al., 2020). First, for each testing night separately, we evaluated the epoch-by-epoch agreement between the human-scoring (considered as the ground-truth) and the algorithm’s predictions. Agreement was measured with widely-used metrics that included accuracy (i.e. percent of correctly classified epochs), Cohen’s kappa (a more robust score that takes into account the possibility of the agreement occurring by chance) and the Matthews correlation coefficient. The latter is thought to be the most robust and informative score for classification as it naturally takes into account imbalance between sleep stages (Chicco and Jurman, 2020). For all of the above metrics, higher values indicate higher accuracy agreement.

Unless specified, we report the performance of the full model which includes one EEG, one EOG, one EMG as well as the age and sex of the participant.

In addition, to measure stage-specific performance, we report confusion matrices and F1-scores for each stage separately. The F1-score is defined as the harmonic mean of precision and sensitivity. Put simply, precision is a measure of the *quality* of the prediction (e.g. the proportion of all the epochs classified as REM sleep by the algorithm that were actually labelled as REM sleep in the human scoring), while sensitivity is a measure of the *quantity* of the prediction (e.g. the proportion of all the epochs labelled as REM sleep in the human scoring that were correctly classified as REM sleep by the algorithm). Being the average of both precision and sensitivity, the F1-score is therefore an optimal measure of the algorithm’s performance that can be calculated independently for each sleep stage. Higher values indicate superior performance.

Additionally, we conducted discrepancy analysis to test for any systematic bias of the algorithm to over- or under-estimate a specific sleep stage. Lastly, we performed moderator analyses to test whether the algorithm maintained a high-level of performance across participants of different ages, gender and health status.

#### Consensus scoring

Each night of the DOD was scored by 5 independent experts. By taking the most voted stage for each epoch, it is therefore possible to build a consensus hypnogram and thus reduce bias caused by low inter-scorer agreement (Guillot et al., 2020; Perslev et al., 2021). When a tie occurs on a specific epoch, the sleep scoring of the most reliable scorer is used as the reference. The most reliable scorer is the scorer with the highest average agreement with all the other scorers. We also compared each human scorer against an unbiased consensus, which was based on the *N-1* remaining scorers (i.e. excluding the current scorer). Here again, the most reliable of the *N-1* scorers was used in case of ties.

### Data and code availability

All polysomnography data can be requested from the NSRR website (http://sleepdata.org). The Dreem Open Dataset can be found at https://github.com/Dreem-Organization/dreem-learning-open. The source code and documentation of the algorithm is available at https://github.com/raphaelvallat/yasa. The Python code to reproduce all the results and figures of this paper can be found at https://github.com/raphaelvallat/yasa_classifier. All analyses were conducted in Python 3.8 using scikit-learn 0.23.2 (Pedregosa et al., 2011) and lightgbm 3.0.0 (Ke et al., 2017).

## Results

### Descriptive statistics

The *training set* consisted of more than 23,000 hours of PSG data across 2,832 unique full-night PSG recordings from seven different datasets (CCSHS, n=417; CFS, n=513; CHAT, n=192; HomePAP, n=72; MESA, n=531; MrOS, n=528; SHHS, n=579). All these datasets are publicly available from the NSRR website (http://sleepdata.org). In total, these 2,832 training nights represent 2,782,081 unique 30-seconds epochs. The demographic characteristics of the training set are as follow: 56% male; age = 51.2 ± 25.4 yrs (median = 59 yrs, range = 5-92 yrs); AHI = 12.38 ± 15.3 (median = 6.74, range = 0-125); BMI =27.71 ± 7.44 (median = 26.6, range = 12.8-84.8); race = 60.5% White, 27% Black, 7.5% Hispanic, 5% Asian or Pacific Islander.

The *testing set 1* consisted of 542 unique full-night PSG recordings from 6 different datasets (CCSHS, n=95; CFS, n=86; CHAT, n=77; MESA, n=93; MrOS, n=91; SHHS, n=100). The testing nights were randomly selected from a subset of nights that were not included in the training set (see **Methods**). We did not include any nights from the HomePAP dataset because the latter only consisted of 72 nights after preprocessing, all of which were used for training. In total, these 542 testing nights represent 541,371 unique 30-seconds epochs, or roughly 4,500 hours of PSG data. The demographic characteristics of the testing set are as follow: 56% male; age = 46.4 ± 27.8 yrs (median = 54 yrs, range = 5-94 yrs); AHI = 12.01 ± 15.5 (median = 6, range = 0-102); BMI = 26.25 ± 7.06 (median = 25.7, range = 13.2-54.3); race = 60.1% White, 28.2% Black, 6.1% Hispanic, 5.5% Asian or Pacific Islander.

The *testing set 2* consisted of 80 full-night PSG recordings from the publicly-available DOD dataset (Guillot et al., 2020). The DOD consists of 25 nights from healthy adults (age = 35.3 ± 7.51 yrs, 76% male, BMI = 23.8 ± 3.4) and 55 nights from patients with obstructive sleep apnea (age = 45.6 ± 16.5 yrs, 64% male, BMI = 29.6 ± 6.4, AHI = 18.5 ± 16.2). Five nights were excluded because the duration of the PSG data and the duration of the hypnogram differed by more than 1 minute, leading to a final sample of 75 nights (~607 hours of data). Each night of the testing set 2 was scored by 5 registered sleep technicians, thus allowing to test the algorithm against a consensus of human scorers (see **Methods**). Unlike the NSRR datasets, which we split into training and testing subsets, the DOD dataset was only used for performance evaluation. This provides a more robust estimation of the algorithm’s performance on real-world data from a new sleep clinic or laboratory.

### Validation results

#### Testing set 1: NSRR holdout

The overall performance of the algorithm on the testing set 1 is described in **Figure 1A**. Median accuracy, calculated across all 542 testing nights, was 85.9%. The median Cohen kappa was 0.791, indicating substantial agreement, and similarly, the median Matthews correlation coefficient was 0.794. The CCSHS database testing set had the highest overall accuracy (median = 88.9%), while the MESA database had the lowest accuracy (median = 82.2%).

**Figure 1:**
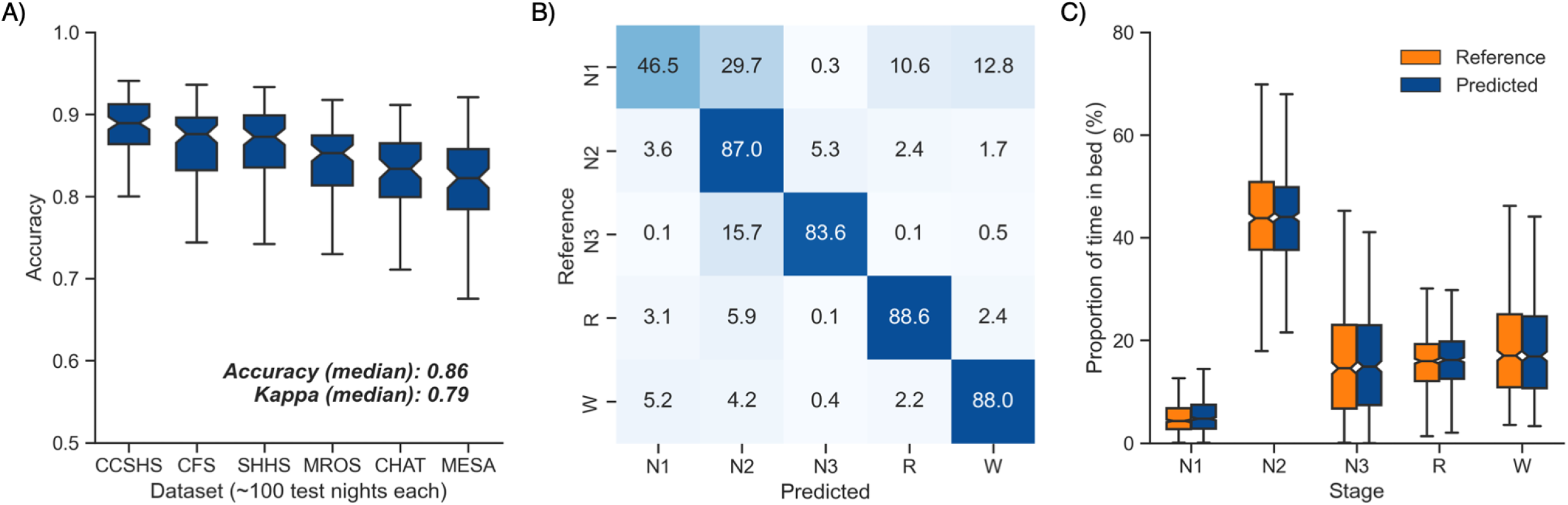
Performance of the algorithm on the testing nights (n=535). **A)** Accuracy of all testing nights, stratified by dataset. The median accuracy across all testing nights was 86%. **B)** Confusion matrix. The diagonal elements represent the percentage of epochs that were correctly classified by the algorithm (also known as sensitivity or recall), whereas the off-diagonal elements show the percentage of epochs that were mislabelled by the algorithm. **C)** Duration of each stage in the human (red) and automatic scoring (green), calculated for each unique testing night and expressed as a proportion of the total time in bed. Error bars represent standard deviations.

Next, we tested the classification performance of individual sleep stages (**Figure 1B**). The overall sensitivity (i.e., the percent of epochs that were correctly classified, see **Methods**) of N3 sleep was 83.6%, leading to a median F1-score across all testing nights of 0.837. REM sleep, N2 sleep and wakefulness all showed sensitivities above 87% (all median F1-scores ≥ 0.87). N1 sleep showed the lowest agreement, with an overall sensitivity of 46.5% and a median F1-score of 0.447 — a finding consistent with this sleep stage often showing very low agreement across human sleep scorers (<45%, (Malhotra et al., 2013)). These algorithm performance results were consistent when examined separately for each database (**Figure S1**).

Further examination of the confusion matrix showed that the two most common inaccuracies of the algorithm were 1) mislabelling N1 sleep as N2 sleep (29.7% of all true N1 sleep epochs) and 2) mislabelling N3 sleep as N2 sleep (15.7% of all true N3 sleep epochs). Importantly, the algorithm made few blatant errors, such as mislabelling N3 sleep as REM sleep (0.1%), or mislabelling REM sleep as N3 sleep (0.06%).

Additional analyses indicated that the algorithm was more prone to inaccuracies around sleep-stage transitions than during stable epochs (**Figure 2F**; mean accuracy around a stage transition: 71.2 ± 6.9, during stable periods: 93.6 ± 5.6%, p<0.001). Similarly, and as expected, accuracy was significantly greater for epochs flagged by the algorithm as high-confidence (≥80% confidence) than in epochs with a confidence below 80% (94.7 ± 4.2% and 62.7 ± 6.4% respectively, p<0.001).

**Figure 2:**
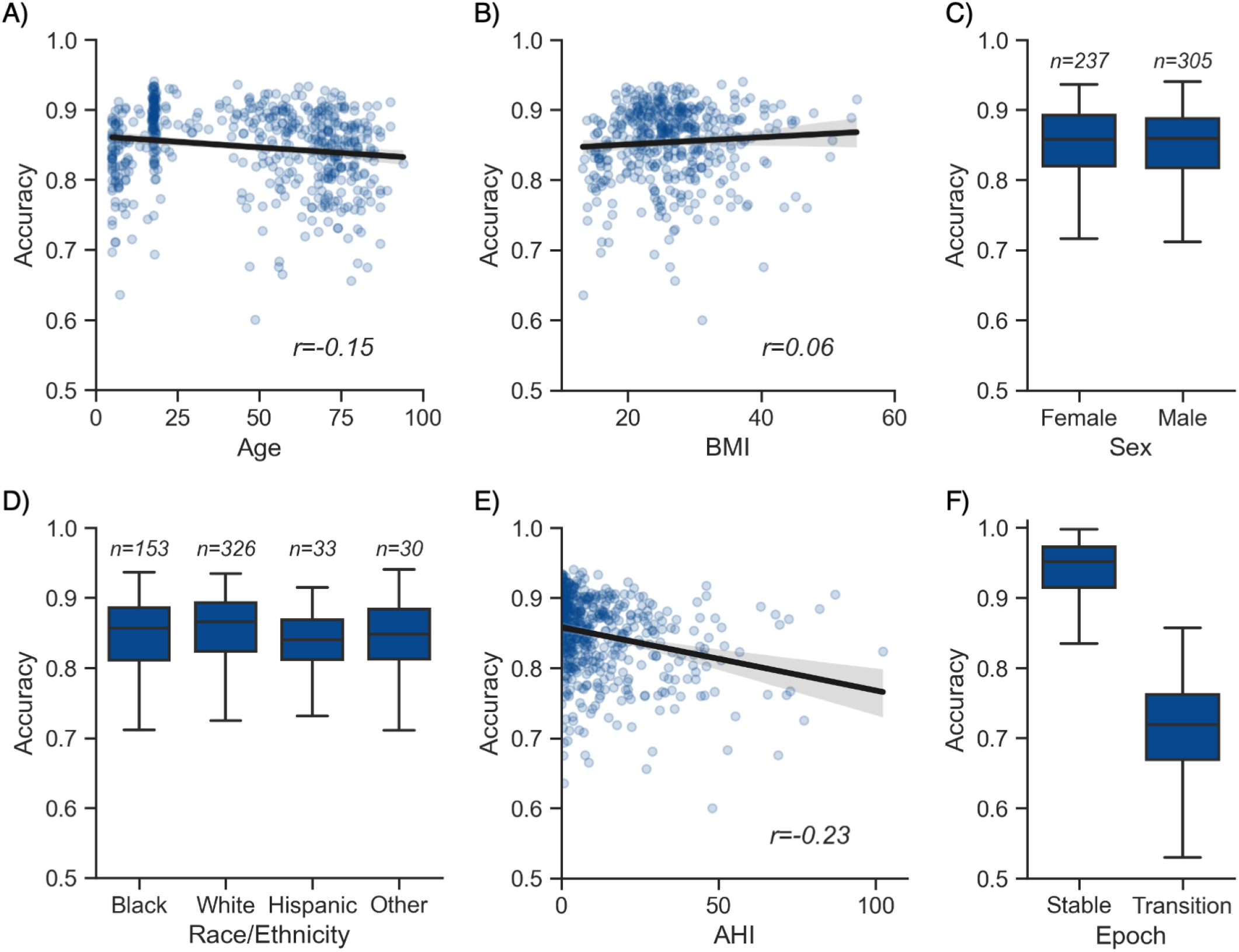
Moderator analyses. Accuracy of the testing nights as a function of age **(A)**, BMI **(B)**, sex **(C)**, race **(D)**, AHI **(E)** and whether or not the epoch is around a stage transition **(F).** An epoch is considered around a transition if a stage transition, as defined by the human scoring, is present within the 3 minutes around the epoch (1.5 minute before, 1.5 minute after).

We further tested for any systematic bias of the algorithm towards one or more sleep stages. The overall proportion of each sleep stage in the testing nights was similar between the human and automatic staging (all Cohen’s d < 0.08; **Figure 1C**), indicating a negligible bias. Similarly, the percentage of stage transitions across the night was consistent in the human and automatic scoring (mean ± STD: 13.5 ± 5.6% and 13.6 ± 4.2% respectively, Cohen’s d=0.03).

#### Testing set 2: consensus scoring on an unseen dataset

Next, we examined the performance of YASA on the testing set 2, a previously unseen dataset of healthy and sleep-disordered breathing patients which was scored by five registered experts (Guillot et al., 2020; Perslev et al., 2021). Median accuracy of YASA against the consensus scoring of the five experts was 85.1%, with a median kappa of 0.770 and a median MCC of 0.774. Median F1-scores for individual sleep stages were 0.368 for N1 sleep, 0.880 for N2 sleep, 0.816 for N3 sleep, 0.903 for REM sleep and 0.846 for Wake. Here again, the performance of the algorithm was significantly higher during stable epochs (90.9 ± 7.3 vs 68.9 ± 7.1, p<0.001) or epochs that were marked as high confidence by the algorithm (93.2 ± 5.9 vs 60.7 ± 7.5, p<0.001).

A comparison of YASA against each of the individual scorers (**Table 1**) revealed that YASA had a significantly better accuracy than 4 out of five experts (two-sided paired T-test: all ps≤0.046), and was not significantly different from the remaining most accurate expert (YASA: 83.3 ± 6.9, best human: 83.3 ± 8.1, p=0.962). A similar pattern was observed with the Cohen kappa: the algorithm was significantly better than 4 out of the five experts (all ps≤0.025), and matched the performance of the best expert (74.7 ± 10.2 vs 74.9 ± 11.7, p=0.916; see **Table 1**).

**Table 1:** Performance of YASA against the five human experts in the DOD dataset (n=75 nights). Mean ± standard deviation scores computed for YASA versus the consensus of all human scorers (last row), or a single human expert versus the consensus scoring of the 4 remaining scorers. Highest scores are highlighted in bold. For similar comparisons of recently-developed algorithms on the same dataset, see (Guillot et al., 2020; Perslev et al., 2021).

|  | Accuracy | Kappa | F1 N1 | F1 N2 | F1 N3 | F1 REM | F1 WAKE |
| --- | --- | --- | --- | --- | --- | --- | --- |
| **Expert 1** | $80.5 \pm 11.9$ | $70.7 \pm 16.7$ | $39.5 \pm 15.2$ | $82.0 \pm 13.4$ | $63.5 \pm 31.5$ | $82.2 \pm 20.8$ | $85.1 \pm 11.8$ |
| **Expert 2** | $80.7 \pm 9.8$ | $71.5 \pm 13.4$ | $44.6 \pm 17.4$ | $82.3 \pm 11.8$ | $63.4 \pm 29.0$ | **$89.1 \pm 12.5$** | $85.1 \pm 11.9$ |
| **Expert 3** | $81.4 \pm 8.7$ | $71.5 \pm 13.1$ | $42.5 \pm 16.1$ | $82.9 \pm 11.0$ | $54.4 \pm 33.0$ | $88.8 \pm 12.8$ | $86.0 \pm 10.8$ |
| **Expert 4** | $79.5 \pm 8.8$ | $69.3 \pm 12.9$ | $39.1 \pm 15.2$ | $82.9 \pm 7.4$ | $60.7 \pm 30.6$ | $86.5 \pm 16.4$ | $82.4 \pm 15.7$ |
| **Expert 5** | $83.3 \pm 8.1$ | **$74.9 \pm 11.7$** | **$46.7 \pm 14.2$** | $85.3 \pm 8.7$ | $66.7 \pm 30.5$ | $86.7 \pm 17.4$ | **$86.3 \pm 11.8$** |
| **YASA** | **$83.3 \pm 6.9$** | $74.7 \pm 10.2$ | $36.5 \pm 17.2$ | **$86.1 \pm 7.2$** | **$71.4 \pm 26.7$** | $87.1 \pm 13.6$ | $81.7 \pm 11.3$ |

In summary, these data demonstrate high levels of sleep-staging accuracy of the algorithm that is on par with or exceeds that of human interscorer agreement and error. Similar to human scoring, the most frequent limitations of the algorithm occured around transition epochs and/or in N1 sleep, while all the other sleep stages showed excellent classification performance. Finally, the algorithm did not systematically under or over-estimate any sleep stages. Instead, the algorithm was successful in preserving the overall distribution of sleep stages across the night.

### Moderator analyses

We further tested the impact of different moderators on the accuracy of the NSRR testing nights, for which extensive demographic data was available. As with human sleep scoring, analyses focused on the first factor of age revealed a small but significant negative correlation with accuracy (r=-0.149, p<0.001, **Figure 2A**), suggesting a moderate linear decrease of accuracy with age. For example, the average accuracy was 87.5% in the 10-20 years old group and 82.4% in the ≥75 years old group. Second, body composition was not significantly correlated with accuracy (r=0.062, p=0.19, **Figure 2B**). Third, sex was not a significant determinant of accuracy either (Welch’s T=0.433, p=0.665, **Figure 2C**). Fourth, sleep apnea, and specifically the apnea-hypopnea index (AHI), was significantly correlated with accuracy (r=-0.23, p<0.001, **Figure 2E**). This would indicate that, to a modest degree, the performance of the algorithm can decline as the severity of sleep apnea increases, possibly mediated by an increased number of stage transitions in patients with sleep apnea (see above stage transition findings). Consistent with the latter, the percentage of (human-defined) stage transitions was significantly correlated with the AHI (r=0.37, p<0.001). Finally, race was not a significant predictor of accuracy (ANOVA, F(3, 538)=0.877, p=0.453, **Figure 2D**).

### Features importance

The 20 most important features of the model are shown in **Figure S2**. Importance was measured on a random sample of 25% of the training set using Shapley values (SHAP; (Lundberg et al., 2020)). Ten out of these top 20 features are derived from the EEG signal, 6 from the EOG and 3 from the EMG. This indicates that EEG is the most important signal for accurate sleep staging, followed by EOG and to a lesser extent EMG. Age is also present in the top 20 features, a result consistent with the well-known changes in sleep stages across the lifespan (Ohayon et al., 2004). Consistent with these findings, the algorithm retained high levels of accuracy when using only a single-channel EEG (median accuracy across the 542 nights of the testing set 1 = 83.5%, median kappa = 0.755) or a combination of one EEG and one EOG (median accuracy = 84.8%, median kappa = 0.779). In other words, using only one EEG and one EOG led to a ~1% decrease in med compared to a full model including one EEG, one EOG, one EMG as well as age and sex (median accuracy = 85.9%, kappa = 0.791).

The single most important feature of the classifier was the absolute power of the EEG power spectrum. This was followed by the EEG beta power, EOG absolute power, and EEG delta/beta ratio. It is relevant that 7 out of the 20 features are relatively uncommon, non-linear features (i.e. Higuchi fractal dimension, Petrosian fractal dimension, permutation entropy). These novel features may capture unique electrophysiological properties of each sleep stage that are missed by traditional linear and/or spectral features. Similarly, seven out of the 20 most important features include some form of temporal smoothing / normalization (e.g. centered 15 minutes rolling average). This highlights the importance of taking into account the neighboring epochs (past and future) for accurate sleep staging.

### Software implementation

Implementation of the algorithm is completely open-source and freely available. Our sleep algorithm, colloquially termed YASA^1^, is part of a broader sleep analysis package (or “library”), written in Python. In addition to the automatic sleep staging module we describe here, YASA also includes several additional functions such as automatic detection of sleep spindles and slow-waves, automatic artifact rejection, calculation of sleep statistics from an hypnogram, spectral power estimation (e.g. **Figure 3B**), and phase-amplitude coupling. However, use of the basic sleep staging module is not at all contingent on a desire to quantify any of these metrics. Simply that they are on offer as additional tools in the software suit, should the user wish.

**Figure 3:**
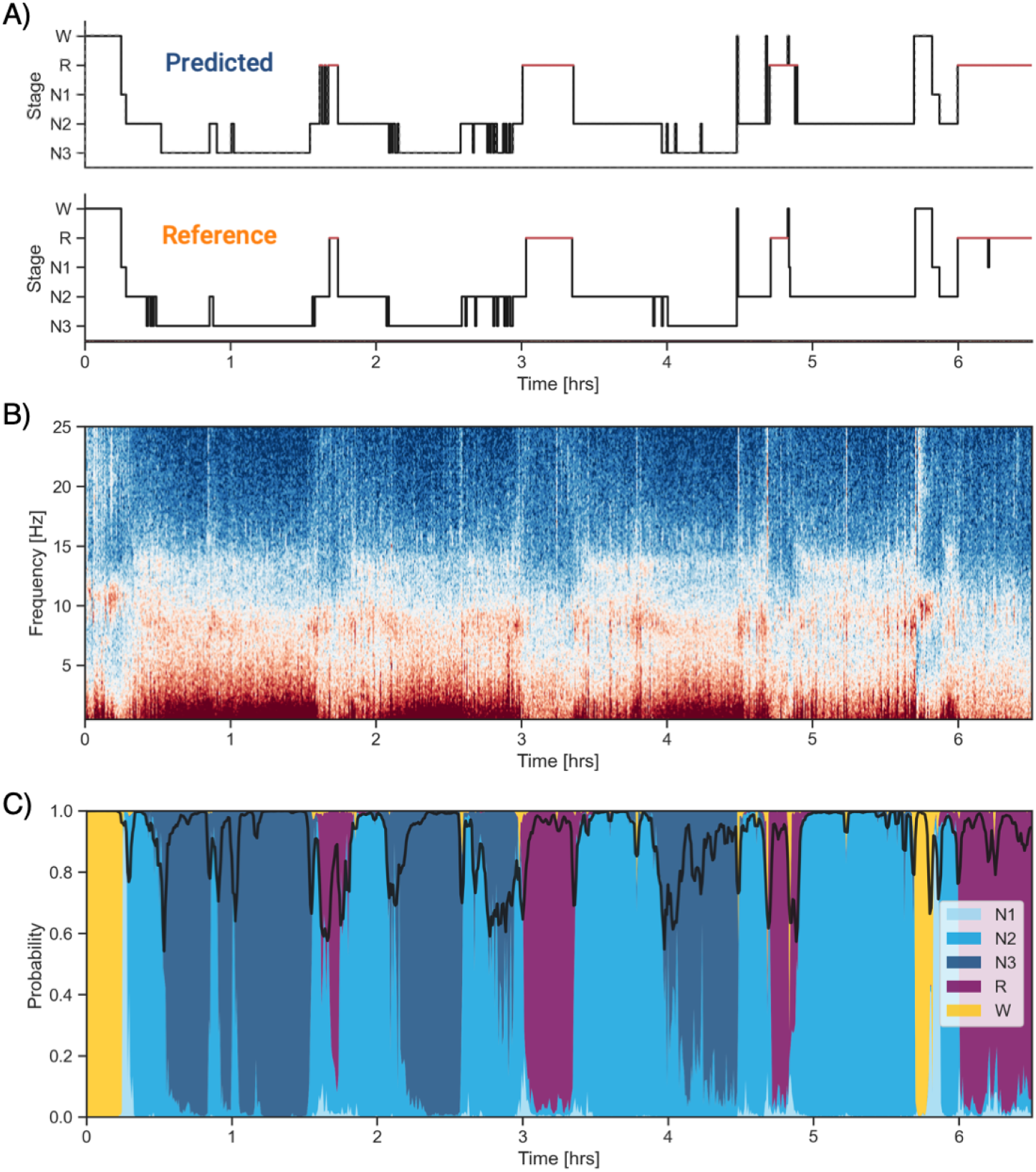
Example of data and sleep stages prediction in one subject. **A)** Predicted and ground-truth (= human-scored) hypnogram in an healthy young female (CFS dataset, 24 yrs old, Black, AHI < 1). The agreement between the two scoring is 92.9%. **B)** Corresponding full-night spectrogram of the central EEG channel. Warmer colors indicate higher power in these frequencies. This type of plot can be used to easily identify the overall sleep architecture. For example, periods with high power in frequencies below 5 Hz most likely indicate deep NREM sleep. **C)** Algorithm’s predicted cumulative probabilities of each sleep stage at each 30-seconds epoch. The black line indicates the confidence level of the algorithm. Note that all the plots in this figure can be readily plotted in the software.

YASA comes with extensive documentation and is released under a BSD-3 Clause license, part of the Open Source Initiative, and can be directly installed with one simple line of code from the Python Package Index repository^2^. The source code of YASA is freely and publicly hosted on GitHub, and follows standard release cycles with a version-tracking of all the changes made to the code and/or pre-trained classifiers. That is, users can choose to ignore the most recent versions and keep a specific version of the code and staging algorithm, which is useful for example in longitudinal studies where the preprocessing and analysis steps should stay consistent across time.

The general workflow to perform the automatic sleep staging is described below (see code snippet in **Figure S3**). First, the user loads the PSG data into Python. Assuming that the PSG data are stored in the gold-standard European Data Format (EDF), this can be done in one line of code using the MNE package (Gramfort et al., 2014), which has a dedicated function to load EDF files^3^. Second, the automatic sleep staging is performed using the algorithm’s sleep staging module^4^. The only requirement is that the user specify the name of the EEG channel they want to apply the detection (preferentially a central derivation such as C4-M1). The algorithm is now ready to stage the data. Should the user wish, there is the option of naming EOG and EMG channels (preferentially a chin EMG). The user can also include ancillary if desired, such as age and sex, which can aid in improving the accuracy of the algorithm, though this is not at all necessary for high accuracy (as described earlier).

Regarding the processing steps of the algorithm, sleep staging is performed with the ‘predict’ function (**Figure 3A**). This function automatically identifies and loads the pre-trained classifier corresponding to the combination of sensors / metadata provided by the user. While pre-trained classifiers are natively included in the algorithm, a user can define custom pre-trained classifiers if they wish, which can be tailored to their specific use cases. Once again, however, this is just for flexibility of use, and is not at all required for the algorithm to perform with high accuracy. Parenthetically, this flexibility can also be leveraged for a variety of use cases, for example, scoring rodent sleep data instead of human sleep data. In addition to stage scoring, implementation of the ‘predict_proba’ function provides the ability for a user to quantify the probability certainty of each stage at each epoch. This probability certainty can then be used to derive a confidence score, should that be desired, although again this is not necessary for standard sleep staging by a user (**Figure 3C**). Finally, the predicted hypnogram and stage probabilities can easily be exported into a text or CSV file using standard Python functions, should the user wish, though once again is not required.

One limitation of automated data analysis methods is that they are computationally demanding (Fiorillo et al., 2019), often requiring specific higher-end computer systems that are costly. To avoid this limitation, the current algorithm is designed to be fast and highly memory-effective by leveraging high-performance tools in Python (e.g. Numpy’s vectorization and Numba compiling). As a consequence, a full-night PSG recording sampled at 100 Hz is typically processed in less than 5-seconds on a rudimentary, basic consumer-level laptop.

## Discussion

Here, we sought to develop a sleep staging algorithm that 1) matches human-scorer accuracy, 2) has been trained on a large and heterogeneous data set, 3) is easy to implement by most individuals, 4) is computational low-demanding and can therefore be run on a basic laptop, and 5) is entirely free, and thus easily adopted by researchers, clinicians and commercial ventures.

### Performance

The algorithm displayed high-levels of accuracy, outperforming those observed across human inter-rater agreement (Fiorillo et al., 2019). First, the median accuracy of the algorithm across 542 testing nights from the NSRR database was 85.9%. Second, on a previously unseen dataset of 25 healthy individuals and 50 patients with OSA, YASA had a significantly better accuracy than 4 out of five clinical experts. The median accuracy of the algorithm against the consensus of five experts was 85.1%. These results match recently developed deep-learning-based algorithms for automatic sleep staging (Guillot et al., 2020; Perslev et al., 2021). Moreover, in both testing sets, the agreement between the algorithm and human scoring was superior to 90% when only considering epochs that occurred during a stable period of sleep (i.e. not around a stage transition).

Regarding individual sleep stages, the algorithm showed excellent classification performance for N2 sleep, N3 sleep, REM sleep and wakefulness, and moderate agreement for N1 sleep. These results are consistent with human inter-rater agreement, which has been found to be consistently lower in N1 sleep compared to the other sleep stages (Malhotra et al., 2013; Norman et al., 2000; Rosenberg and Van Hout, 2013). Furthermore, the algorithm was successful in preserving the overall distribution of sleep stages across the night, such that is neither over- or under-estimating a specific sleep stage.

Beyond a basic sleep stage classification, an advantage of the algorithm is its ability to provide probability (i.e. likelihood) value for each individual epoch of each stage. These probabilities inform the users about the confidence level of the algorithm. As such, the algorithm provides a unique feature addition for sleep scoring that extends beyond sleep-staging quantification, and layers atop sleep-staging qualitative assessment. Proving the validity of this measure, the accuracy of the algorithm was statistically superior in epochs that were flagged as high-confidence, reaching ~95% agreement with human scoring. Among many other advantages, such an ability could be used in a semi-supervised scoring approach if desired, wherein a human scorer can focus expressly on the selection of low-confidence epochs for attention, limiting time investment.

### Generalizability

The power and utility of an algorithm is not just determined by its accuracy, but also by the characteristics of the underlying dataset that was used for training and validation (Fiorillo et al., 2019). For example, an algorithm showing excellent performance but that was trained on a very small and heterogeneous sample size (e.g. 10-20 healthy young adults from the same demographic group and recorded using the same device) will typically have low utility for other use cases, such as scoring data from another PSG device, or data from patients with sleep disorders. The generalizability of such algorithms can therefore be compromised, with the high-performance most likely indicating over-fitting to the small training dataset.

Seeking to avoid this issue, our algorithm was trained and validated on more than +27,000 hours of PSG human sleep recordings across +3000 nights, from numerous independent and heterogeneous datasets. These datasets included participants from a wide age range, different geographical locations, racial groups, sex, body composition and health status. Importantly, these independent studies involved different recording devices and settings (e.g. montage, sampling rate). Such a high heterogeneity is paramount to ensure a high reliability and generalizability of the algorithm.

Moderator analyses showed that the performance of the algorithm was unaffected by race, sex and body composition. Consistent with human scoring (Muehlroth and Werkle-Bergner, 2020; Norman et al., 2000), the performance of the algorithm was sensitive to advancing age and increasing severity of sleep apnea, although accuracy remained high (>80%). The latter two circumstances can be explained by the increase in the number of stage transitions and the increase in the proportion of N1 sleep typically seen in aging and sleep apnea (Norman et al., 2000; Ohayon et al., 2004). Indeed, the majority of such scoring errors were located around stage transitions and/or in N1 sleep, which is consistent with findings of human inter-rater error of these same circumstances (Norman et al., 2000).

Another problematic challenge in the field of automated sleep staging is the fact that each algorithm is usually tested on a different (often non-publicly available) dataset, which renders the comparison of algorithms nearly impossible. Adding to this, prior approaches are also heterogeneous in the performance evaluation metrics used to validate the algorithm, as well as in the PSG channel(s) on which the algorithm is applied — with some algorithms using only a single EEG channel and others a multi-channel approach (Fiorillo et al., 2019). Addressing these issues, the current algorithm was trained and tested on publicly available datasets, using standard classification metrics (e.g. accuracy, F1-scores, confusion matrix) and following recent guidelines for performance evaluation (Menghini et al., 2020). Furthermore, the built-in flexibility of the algorithm allows for different combinations of channels to be used, should a user wish. For all these reasons we hope that the algorithm is not only of broad utility, but will facilitate replication of our work and comparison to existing and future works.

### Easy of use and computationally low demand

To facilitate wide adoption by the sleep community, it is of utmost importance that any algorithm can be used and understood by all parties concerned (e.g. students, researchers, clinicians, technicians), no matter the level of technical expertise. To ensure this, the software has been built with a particular focus on ease-of-use, documentation and transparency.

First, the end-to-end sleep staging pipeline is written in less than 10 lines of Python code, and the software comes with pre-trained classifiers that are automatically selected based on the combination of channels used, thus limiting the risk of any error. Second, the software has extensive documentation and includes numerous example datasets, allowing any users to get familiarized with the algorithm before applying it to their own datasets, if they wish (though it is not necessary). Third, the algorithm uses a traditional features-based approach to classify sleep stages, instead of a black-box algorithm. These features are described in detail in the documentation and source code of the algorithm, and can be explained to any researchers or clinicians in lay terms. Another advantage of using features-based versus prior deep learning algorithms is that model training is significantly faster and does not require specialized hardware (e.g. high-performing graphics processing unit) — YASA’s model training can be done in less than an hour on a basic laptop. Finally the sleep staging is done locally on the user’s computer, and the data is never uploaded to the cloud or any external servers, thus limiting security and privacy risks (Fiorillo et al., 2019), and the need for any connectivity when using the software.

### Free, open-source and community-driven

Another potential reason that automated sleep staging has yet to become a de facto standard is that some algorithms are sitting behind a paywall (e.g. (Malhotra et al., 2013; Patanaik et al., 2018)). By contrast, the current algorithm is free and released under a non-restrictive BSD-3 open-source license. The software, which also includes other sleep analysis tools (e.g. sleep spindle detection, spectral estimation, automatic artifact rejection, phase-amplitude coupling), is hosted on GitHub and has already been downloaded several thousand times at the date of writing^5^. The software can therefore be used by for- or non-profit outlets for free, without constraint.

### Conclusion

The interest in quantifying and tracking of human sleep has increased exponentially in the past decade (Fleming et al., 2015; Shelgikar et al., 2016), and demand for a highly automated, accurate, high-speed, and easy to implement algorithms for the scoring of human sleep has scaled similarly. Here, we offer an algorithm that seeks to accomplish this need. It is our hope that this free tool has the potential for broad adoption across all outlets (research, clinical, commercial) and use cases. With further validation and feedback from the sleep community, we hope it can become an industry standard method, one that can be built upon, expanded and refined through cross-collaborative open-source community development.

## Abbreviations

AHI: apnea-hypopnea index
BMI: body mass index
EEG: electroencephalogram
EOG: electrooculogram
EMG: electromyogram
OSA: obstructive sleep apnea
PSG: polysomnography
MCC: Matthews correlation coefficient
NREM: non rapid eye movement (sleep)
REM: rapid eye movement (sleep)

## Supplementary Figure Captions

**Figure S1: Performance on the testing nights, stratified by dataset.** The diagonal elements of the confusion matrices represent the percentage of epochs that were correctly classified by the algorithm (also known as sensitivity or recall), whereas the off-diagonal elements show the percentage of epochs that were mislabelled by the algorithm.

**Figure S2: Top 20 most important features of the classifier.** Importance was measured on a random sample of 25% of the training nights during model training using Shapley values (SHAP).

**Figure S3: Code snippet illustrating the simplest usage of the algorithm.** Here, automatic sleep staging is applied on an EDF file, previously loaded using the MNE package. The sleep stages predictions are made using one EEG, one EOG and one EMG as well as the age and sex of the participant. The full hypnogram is then exported into a CSV file, together with the epoch number.

## Footnotes

1 https://github.com/raphaelvallat/yasa

2 In a terminal: pip install yasa

3 https://mne.tools/stable/generated/mne.io.read_raw_edf.html

4 https://raphaelvallat.com/yasa/build/html/generated/yasa.SleepStaging.html

5 https://static.pepy.tech/badge/yasa

